# Early onset of Ca^2+^ waves and synchronization in multicellular clusters facilitate focal arrhythmogenesis in human heart failure

**DOI:** 10.1101/2025.05.02.651991

**Authors:** Darya Kazakova, Michael A. Colman, Ankit Pradhan, Lukas Gudaitis, Luka Nys, Bert Cools, Filip Rega, Bert Vandenberk, Cesare Terracciano, H. Llewelyn Roderick, Karin R. Sipido, Eef Dries

## Abstract

**Background:** Spontaneous Ca^2+^ release events and waves are frequent in isolated ventricular cardiomyocytes from failing hearts (HF) and are proposed to initiate arrhythmias in the intact heart. However, evidence supporting whether single-cell Ca^2+^ waves trigger tissue-wide depolarization in the intact heart is scarce, particularly in human HF. We characterized Ca^2+^ waves at single-cell resolution within the multicellular network of the intact heart and identified propagating dynamics and mechanisms facilitating arrhythmogenesis at tissue level.

**Methods:** Living myocardial slices (LMS) from HF and non-HF human hearts were prepared from left ventricular tissue and paced at 2 Hz under adrenergic stimulation. Ca^2+^ transients and waves were recorded by wide-field imaging of Fluo-8. Ca^2+^ waves in relation to single-cell structures within each LMS were identified using custom algorithms. Computational modelling assessed whether experimentally observed HF Ca^2+^ waves dynamics can lead to focal excitation in tissue models.

**Results:** Following pacing, early onset Ca^2+^ waves, initiating within the first 2 seconds, were more frequent in HF compared to non-HF, and HF cardiomyocytes had more foci, where Ca^2+^ waves originate, than non-HF. Spatial mapping showed that early onset waves in HF occurred frequently in clusters of neighboring cells. Although early onset Ca^2+^ waves propagated similar distances in HF and non-HF cardiomyocytes, they more frequently crossed cell boundaries in HF. Particularly, HF LMS exhibited more side-to-side Ca^2+^ propagation, correlating with increased connexin 43 distribution to lateral membranes. Furthermore, HF LMS exhibited more local and global triggered Ca^2+^ activities compared to non-HF LMS, correlating with local tissue depolarization. Simulations of HF Ca^2+^ wave dynamics in remodeled tissue demonstrated a greater capacity to elicit focal excitation.

**Conclusions:** In human HF, a higher incidence of early onset Ca^2+^ waves combines with altered intercellular connectivity to create synchrony in clusters of nearby cells that can overcome the current sink, thereby increasing arrhythmia susceptibility.

**GRAPHICAL ABSTRACT:** 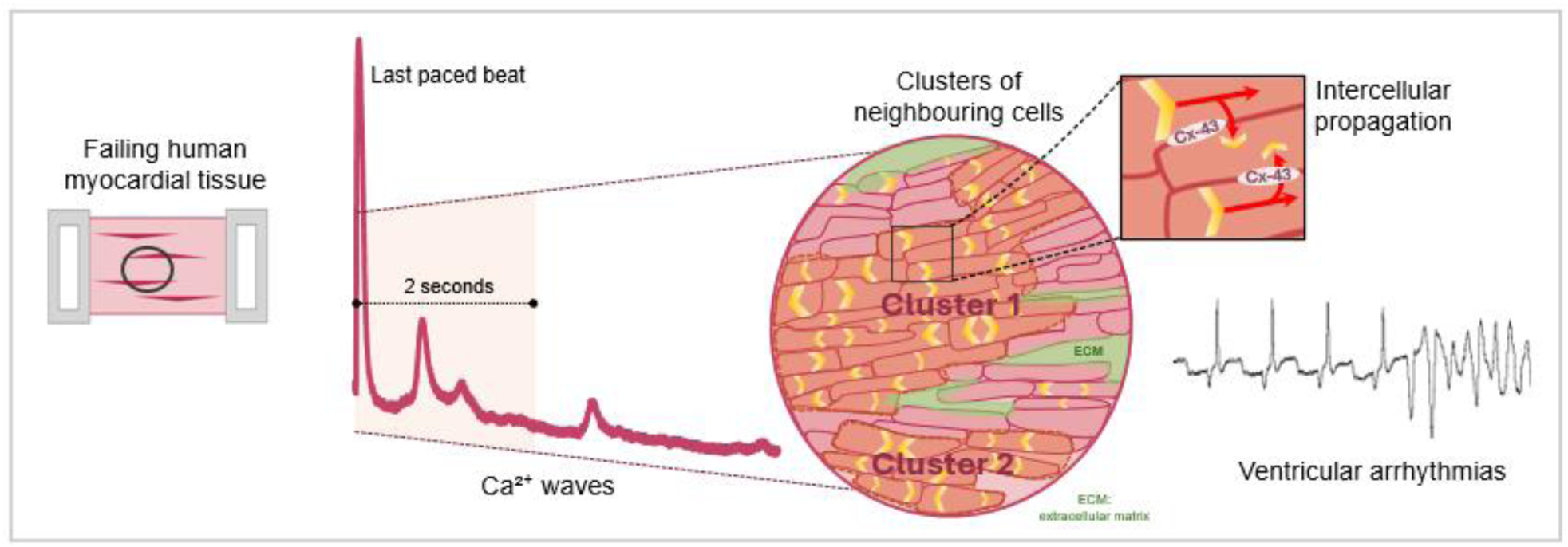

## INTRODUCTION

Arrhythmias are a major cause of death in heart failure (HF) and account for the majority of cases of sudden cardiac death^1^. Unfortunately, treatment options are limited and currently a major role remains for ablation therapy or implantable cardioverter defibrillators, which can terminate a life-threatening arrhythmic event, but do not prevent occurrence^2,3^. Antiarrhythmic drugs have limited applications because of low efficiency and side-effects^3^, though HF medication can indirectly reduce arrhythmic events through reducing cardiac remodelling that underlies mechanisms for arrhythmia induction^4^. Indeed, the failing heart is a vulnerable substrate for arrhythmia with remodeled cardiomyocytes and fibrosis facilitating re-entry, upon which triggers such as premature ventricular complexes (PVCs) can initiate ventricular arrhythmias^5,6^. Modulating events will precipitate PVCs, and a typical example is a surge in adrenergic tone and circulating catecholamines as cause of triggered arrhythmias. Novel avenues for pharmacotherapy in arrhythmic disorders, such as CPVT, target cellular mechanisms of triggers and could provide new options in HF as well, but require more evidence and mechanistic description.

Cardiomyocytes are central actors in generating triggered action potentials (AP) to initiate PVCs. At the cardiomyocyte level, spontaneous Ca^2+^ release from the sarcoplasmic reticulum via ryanodine receptors (RyRs), large enough to become a cell-wide Ca^2+^ wave, generates an inward current through the Na^+^/Ca^2+^ exchanger (NCX), resulting into delayed afterdepolarizations (DADs)^7–10^. If the amplitude of the DAD reaches the threshold for activation of Na^+^ channels, a spontaneous AP is triggered. Such a cellular event can generate a local depolarization in the heart sufficiently large to propagate as a PVC, given a sufficient mass of cardiomyocytes, synchronization and propagation^11–14^. Alternative mechanisms for generating a PVC are local re-entry, which do not rely on spontaneous Ca^2+^ release^5,15,16^.

The synchronization of isolated cellular events is most likely through cell-to-cell conduction of local depolarizing currents, and, as for the eventual propagation throughout the tissues, occurs via gap junctions (GJs). In HF, the major ventricular cardiomyocyte isoform of the GJ protein, connexin43 (Cx43), is redistributed from the cell ends at the intercalated discs (IDs), where it facilitates conduction in a longitudinal direction towards the lateral cell side^17–19^. This lateralization can result in impaired cell-to-cell communication and conduction velocity, contributing to the substrate for re-entry and facilitating arrhythmias^20,21^. Whether the lateralization contributes to triggers for arrhythmia is not established. However, it may reduce electrotonic load, which could otherwise help suppress cellular-scale DADs, potentially facilitating their propagation.

While studies in animal models support the concept that HF cardiomyocytes are more prone to arrhythmogenic Ca^2+^ release events and Ca^2+^ waves^22–25^, data from human hearts is less abundant. Studies on isolated human ventricular cardiomyocytes have reported higher incidence of Ca^2+^ waves^26,27^, and studies at whole heart and tissue level have documented arrhythmia initiation and propagation^28,29^. However, direct evidence for the transition of single-cell events into a tissue-wide depolarization and PVC is scarce. Contributing to this lack of evidence are challenges associated with imaging of arrhythmic events in the same preparation simultaneously at tissue and single-cell resolution. At this time, support for the concept of expansion of events at single-cell or cluster level into tissue-wide events, comes mostly from computational modeling. Limited experimental data are available in animal models, however without reaching high resolution at single-cell level^30,31^, and to the best of our knowledge there are no data for the human heart.

We investigated whether and how Ca^2+^ waves originating within a single cardiomyocyte initiates an AP with tissue-wide propagation in human HF. From a computational point of view, the discrepancy between the source and sink of electrical currents may hinder the spread of spontaneously generated depolarizations from one cell to its adjacent cells^32,33^. Therefore, a propagated signal would require a large mass and synchronization. Whether local Ca^2+^ waves can propagate via GJ and thereby create the local critical mass of cardiomyocytes is in theory possible, but with little experimental evidence. A combination of both electrical and Ca^2+^ propagation is also possible^34^. The general question thus leads to specific parameters to be investigated: the occurrence and kinetics of Ca^2+^ waves in single cardiomyocytes within a connected tissue, the propagation of these events, and whether they can be synchronized and induce a tissue-wide event.

We take advantage of recent methodological developments, and use human living myocardial slices (LMS), thin sections of the human heart where unique features of the adult myocardium are preserved. In the preparation, we can image Ca^2+^ waves in single cells, as well the tissue-wide behavior. As such, LMS offer a new platform to study mechanisms that lead to arrhythmias at the tissue level. They also provide a direct translational entry point to allow the study of human tissue from HF in comparison to tissue from non-failing hearts.

## MATERIAL AND METHODS

Methods are briefly described here. A detailed description of the methods are provided in Supplement. Left ventricular (LV) tissue samples for LMS preparation were obtained from explanted human failing (N=12) and non-failing non-used donor hearts (N=14) (**Figure 1A, *Table S1***). LMS preparation followed an established protocol^35,36^ (**Figure 1B**). LMS were incubated with the Ca^2+^ dye Fluo8-AM and voltage sensitive dye Di-4-ANBDQPQ in normal Tyrode solution, containing 0.1 % Pluronic**®** F-127 and 20 µM (-)-Blebbistatin. After incubation, LMS were transferred to an upright Zeiss AxioExaminer microscope equipped with an LED light source, a rapid acquisition camera and a 10× water-dipping objective, with constant perfusion of normal Tyrode solution at 37°C, containing 15 μM (-)-Blebbistatin. To induce cellular Ca^2+^ waves, LMS were paced for 2 min using electrical field-stimulation at 2 Hz, in the presence of isoproterenol (ISO, 50 nM). After 2 min, pacing was stopped and Ca^2+^ waves were recorded. The temporal and spatial Ca^2+^ waves data were analysed in the 20 s after the last paced beat using TrackMate^37^ with the StarDist detector^38^ and linear assignment problem tracker (**Figure 1C**). The parameters for automated tracking were defined and validated against manual tracking (***Figure S1*, *Video S1-S2***). Frozen OCT samples were used to prepare cryosections and stained with primary antibodies for simultaneous visualization of ID (mouse anti-β-Catenin) and Cx43 (rabbit anti-Cx43). Secondary antibodies, along with Alexa-Fluor647-conjugated wheat germ agglutinin (WGA) to report on cell membranes, were applied to stain Cx43 and anti-β-Catenin using Alexa-Fluor568 (donkey anti-rabbit) and Alexa-Fluor488 (goat anti-mouse), respectively. Images were acquired using a confocal Nikon Ti2 microscope and the percentage of lateral Cx43 labeling was quantified using CellProfilerv4.2.1. Computational modeling was used to assess whether experimentally observed Ca^2+^ waves dynamics can lead to focal excitation in tissue models. A phenomenological model of spontaneous Ca^2+^ release^34,39,40^ was used, of which the “spontaneous release function” (SRF) approach^34^ is ideal for direct parameterisation by the measurements performed in this study. A detailed description of the models are in the ***Supplemental Material***, ***Table S2*** and ***Figures S2-S4***.

**Figure 1.**
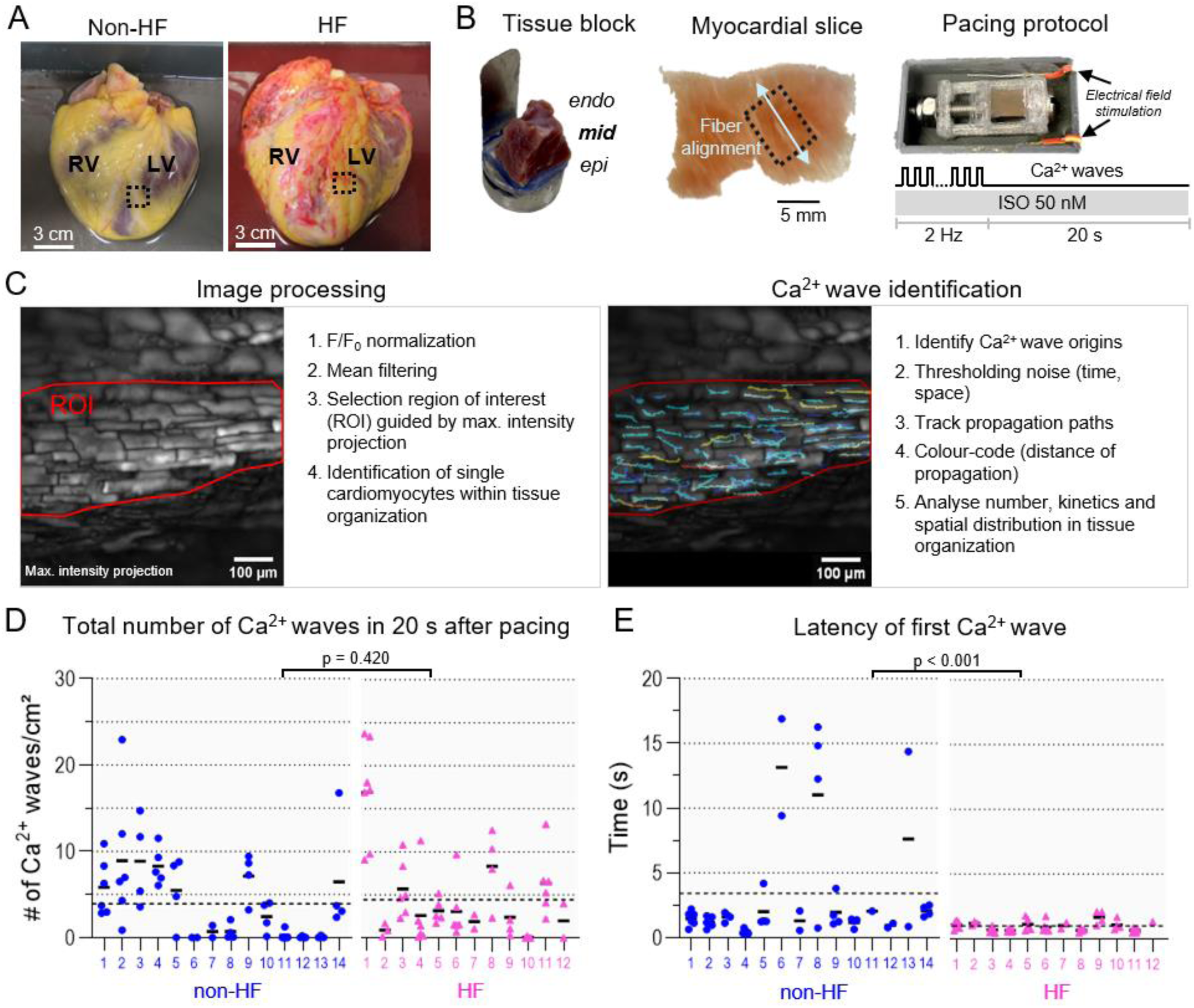
The initiation of Ca^2+^ waves starts early after pacing in HF. **(A)** Example images of human non-failing (non-HF) and failing (HF) hearts. Left ventricle (LV), right ventricle (RV) and regions where transmural tissue blocks were taken (box) are indicated. **(B)** Living myocardial slices (LMS) were prepared from LV transmural tissue blocks. A region with longitudinal fiber alignment was selected from mid-myocardium LMS. LMS were electrically field-stimulated in presence of isoproterenol (ISO). Ca^2+^ waves were recorded for 20 s after the last electrically-triggered Ca^2+^ transient. **(C)** Single-cell Ca^2+^ waves were identified within a region of interest (ROI) and wave kinetics and spatial distribution were analysed. **(D)** Total number of Ca^2+^ waves in 20 s at rest. Mean values ± SEM: 3.94 ± 0.97 s in non-HF (n_LMS_=57) vs. 4.44 ± 1.32 s in HF (n_LMS_=55). **(E)** The latency to the first Ca^2+^ wave. Mean values ± SEM: 3.43 ± 1.09 s in non-HF (n_LMS_=50) vs. 1.01 ± 0.09 s in HF (n_LMS_=54). (Each dot represents one LMS, with mean per heart (black line) and across hearts (dashed line). Individual hearts are indicated by number, N=14 non-HF; N=12 HF).

### Statistics

For normally distributed data, comparisons were made using a Student’s t-test and a hierarchical nested t-test, where appropriate. For non-normally distributed data, either a log-transformation was applied to achieve normality, comparisons were made using the Mann-Whitney test or a linear mixed-model was used to account for repeated measures and hierarchical structure. Gaussian fitting was used for curve fitting and comparisons were made for means, standard deviations (SDs), and amplitudes. Categorical data were assessed using Fisher’s exact test. Data was considered statistically significantly different when p-value < 0.05.

## RESULTS

### Ca^2+^ waves initiate early after paced beats in HF

Single cardiomyocytes within the tissue organization were identified in the images of maximal intensity projection obtained by overlaying all Ca^2+^ waves after pacing was stopped (**Figure 1C, left**). Within this organizational framework, the cellular origin of Ca^2+^ waves could be identified, and the waves’ kinetics and their spatial distribution could be analyzed (**Figure 1C, right**). Somewhat unexpectedly, the total number of Ca^2+^ waves recorded in 20 s was not significantly different between non-HF and HF LMS (**Figure 1D**). However, the latency of the first Ca^2+^ wave (i.e. the time to the initiation of the first Ca^2+^ wave) was significantly shorter in HF LMS compared to non-HF LMS (**Figure 1E**).

### Ca^2+^ waves with short latency are more frequent in HF

Next, we dissected the temporal distribution of Ca^2+^ waves within each LMS. We constructed distribution maps of the sites of Ca^2+^ wave initiation, showing the latency of each Ca^2+^ wave origin in relation to cardiomyocyte structures. From these maps, the latency for each Ca^2+^ wave was extracted and values were categorized into 5 s time bins (**Figure 2A**). Except for one HF and two non-HF LMS, more than 30% of all Ca^2+^ waves occurred in the first 5 s after pacing (**Figure 2B**). We further dissected the distribution of Ca^2+^ waves in this 5 s window by categorizing events into 500 ms bins (**Figure 2C**). Curve fittings of the distribution plots showed a significant leftward shift in the timing of the first Ca^2+^ wave after pacing in HF compared to non-HF, confirming the short latency (i.e. reduced time to initiation) of individual Ca^2+^ waves in HF. This leftward shift was most prominent below 2 s (**Figure 2C**), which falls within the diastolic time intervals of the working myocardium. Therefore, we examined whether early onset Ca^2+^ waves are present in diastolic intervals during pacing. We used 1 Hz pacing, allowing for sufficient diastolic time to record inter-beat Ca^2+^ waves, and compared pacing with and without ISO (**Figure 2D, *Video S3-4***). In HF LMS, more inter-beat Ca^2+^ waves were observed compared to non-HF LMS, but only in the presence of ISO (**Figure 2E**). Furthermore, early onset Ca^2+^ waves in synchrony generated larger amplitudes in HF compared to non-HF (***Figure S5***).

**Figure 2.**
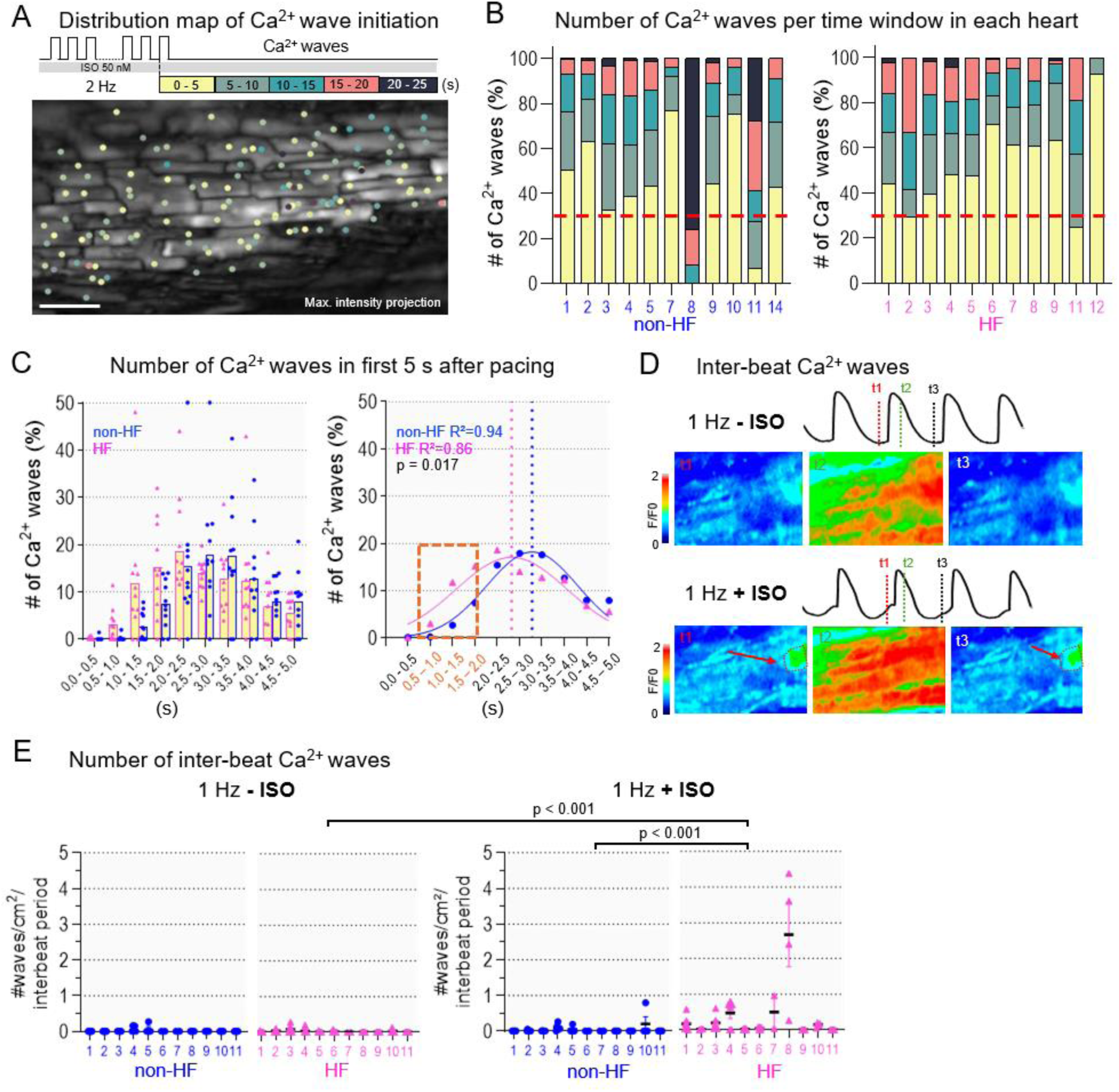
HF has more Ca^2+^ waves with reduced latency. **(A)** Example map of the time to initiate Ca^2+^ waves (latency) in LMS. Each dots represents one Ca^2+^ wave origin. Latency time is shown in bins with different color codes. Scale bar equals 100 µm. **(B)** Stacked bar graphs for each heart with the number of Ca^2+^ waves per time window in non-HF and HF LMS. **(C)** Left: distribution of the number of Ca^2+^ waves with different initiation times in non-HF and HF LMS in the first 5 s at rest. Right: Gaussian curve fits of the distributions in the left panel and comparison between non-HF and HF curve fits. Dotted lines show the mean ± SD: 3.28 ± 1.02 s in non-HF; 2.82 ± 1.24 s in HF; N=11 non-HF; N=11 HF. **(D)** Example images and traces of inter-beat Ca^2+^ waves at 1 Hz with and without isoproterenol (ISO). Different time frames are indicated with t_x_. **(E)** The number of inter-beat Ca^2+^ waves in non-HF vs. HF LMS with and without ISO. Mean values ± SEM: 0.02 ± 0.01 waves/cm²/interbeat period in non-HF without ISO (n_LMS_=46); 0.02 ± 0.01 waves/cm²/interbeat period in HF without ISO (n_LMS_=53); 0.04 ± 0.02 waves/cm²/interbeat period in non-HF with ISO (n_LMS_=46); 0.38 ± 0.13 waves/cm²/interbeat period in HF with ISO (n_LMS_=53). (Each dot represents one LMS, with mean per heart (black line). Individual hearts are indicated by number, N=11 non-HF; N=11 HF).

### Early onset Ca^2+^ waves in HF occur in close proximity

The likelihood of inducing arrhythmogenic activity depends on both timing and proximity of Ca^2+^ waves to each other. We investigated the spatial distribution of early onset Ca^2+^ waves by measuring the distance from the origin of each Ca^2+^ wave to the nearest neighbor Ca^2+^ wave (NND) (**Figure 3A**). Ca^2+^ waves were categorized into 5-µm NND bins; most events were included within a NND of 40 µm (**Figure 3B**). Curve fittings of the distribution plots showed a significant leftward shift in HF compared to non-HF (**Figure 3B, right**), indicating that in HF Ca^2+^ wave origins are closer to each other compared to in non-HF. Moreover, in HF, cells with early onset Ca^2+^ waves appear to cluster in groups, defined as ≥ 5 neighboring cells with early onset Ca^2+^ waves (**Figure 3C**). More of these clusters appear in HF LMS (**Figure 3D**), and these clusters contain more cells compared to non-HF (**Figure 3E**). Furthermore, at the level of single cells, more Ca^2+^ wave origins per cell were observed in HF compared to non-HF LMS (**Figure 3F**). Together, these data indicate that early onset waves occur in close proximity in HF LMS, with multi-origin cells arranged closely together.

**Figure 3.**
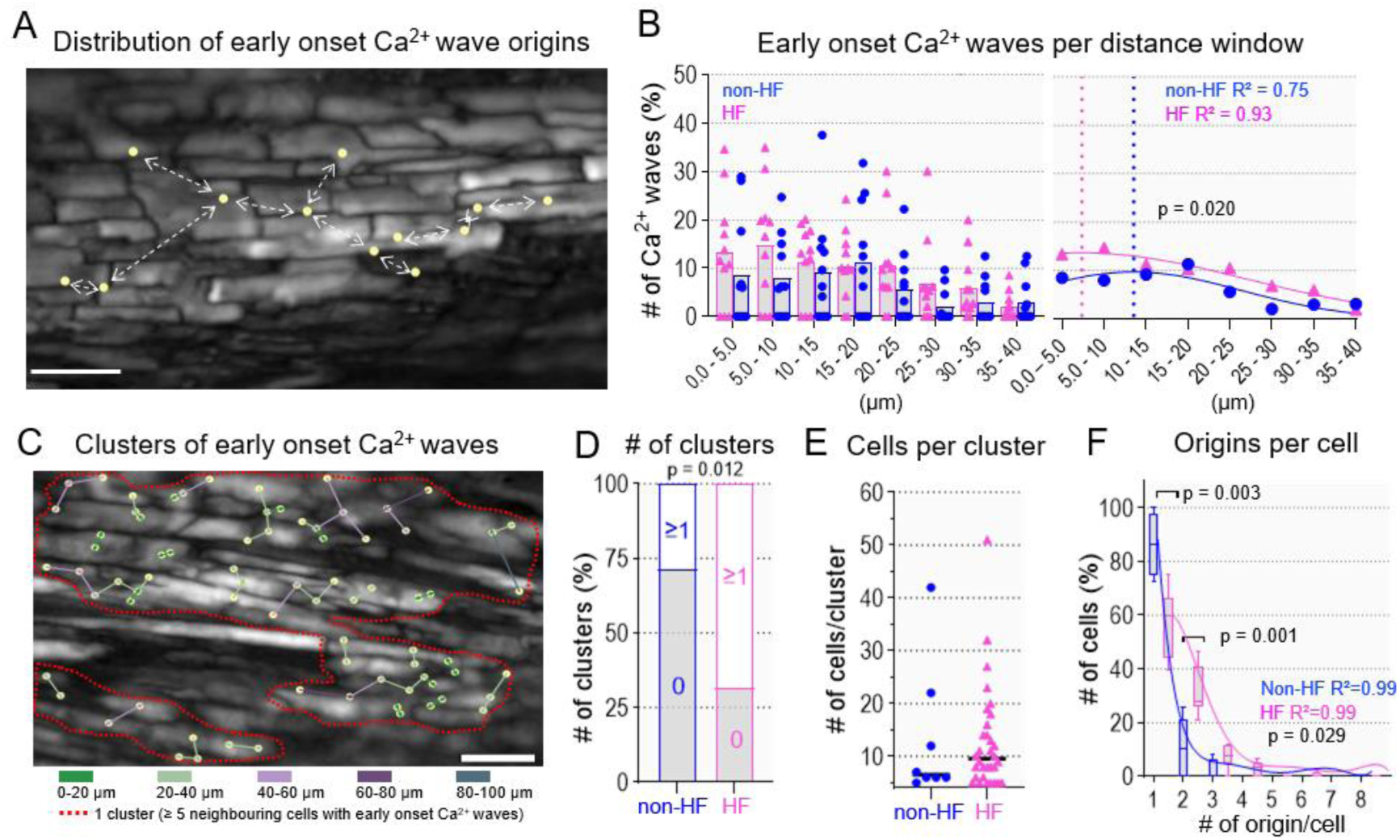
Ca^2+^ waves with early onset occur in close proximity in HF. **(A)** Example map of nearest neighboring distance (NND) between early onset Ca^2+^ waves in LMS. Each dot represents an individual Ca^2+^ wave origin, the arrows depict NND. Scale bar equals 100 µm. **(B)** Left: distribution of the number of waves with different NND in non-HF and HF LMS in the first 2 s at rest. Right: Gaussian curve fits of the distributions in the left panel and comparison between non-HF and HF curve fits. Dotted lines show the mean ± SD: 13.55 ± 12.85 µm in non-HF; 7.37 ± 19.58 µm in HF; N=11 non-HF; N=11 HF. **(C)** Example map of clusters (red) with early onset waves. Each dot represents an individual Ca^2+^ wave origin, color-coded arrows depict NND distance. **(D)** Fraction of LMS with clusters in non-HF vs. HF (8/28 LMS in non-HF and 26/38 LMS in HF). **(E)** Number of cells per cluster in non-HF (median [IQR]: 6.5 [6.0 to 19.5] cells/cluster, N=8) vs. HF (median [IQR]: 9.5 [6.0 to 15.5] cells/cluster, N=32). **(F)** Distribution and curve fits of the number of Ca^2+^ wave origins per cell in non-HF vs. HF measured 2 s after the last pacing beat.

### Intercellular propagation of Ca^2+^ waves in HF

In HF, the average propagation velocity of early onset Ca^2+^ waves was significantly higher compared to non-HF (**Figure 4A**), while the propagation distance was comparable in HF and non-HF. Moreover, most of the Ca^2+^ waves did not propagate beyond 100 µm, suggesting that they remain within cardiomyocytes (**Figure 4B**). However, when we examined wave progress across cells borders, we detected intercellular propagation of Ca^2+^ waves (**Figure 4C**, ***Video S5-6***) in 55% and 32% of the LMS in HF and non-HF, respectively (**Figure 4D**). Compared to the total number of Ca^2+^ waves, the Ca^2+^ waves that propagated intercellularly made up 4.0% and 1.7% in HF and non-HF, respectively (**Figure 4E**). In HF, intercellular Ca^2+^ waves predominantly propagated in a side-to-side direction (59% of intercellular Ca^2+^ waves), a pattern not commonly seen in non-HF (24% of intercellular Ca^2+^ waves), where most intercellular waves propagated from cell end to end (**Figure 4F**). Regardless of the condition, the propagation path of intercellular Ca^2+^ waves involved mainly two neighboring cells but in some cases three (**Figure 4G**). Propagation into four cells was observed only once in HF (**Figure 4G**).

**Figure 4.**
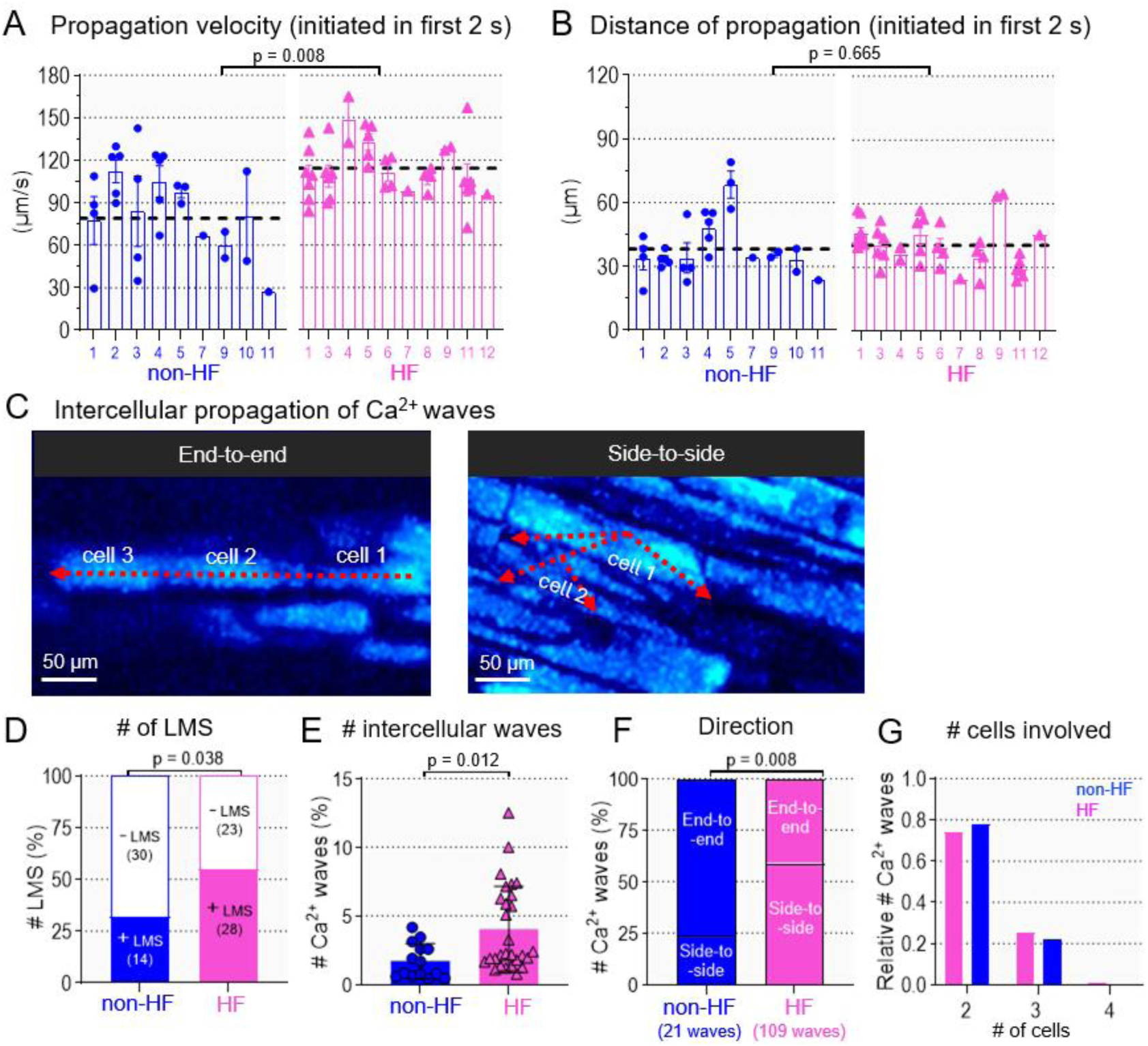
Kinetics and intercellular propagation of Ca^2+^ waves. (A) Velocity of early onset Ca^2+^ waves. Mean values ± SEM: 78.9 ± 8.62 µm/s in non-HF (n_LMS_=27; N=9); 114.3 ± 5.25 µm_/s_ in HF (n_LMS_=38; N=10). (B) Propagation distance of early onset Ca^2+^ waves. Mean values ± SEM: 38.2 _± 4_.31 µm in non-HF (n_LMS=2_7; N=9); 40.4±3.43 µm in HF (n_LMS_=38; N=10). (C) Left: Example images of intercellular propagation in end-to-end or side-to-side mode. (D) Fraction of LMS with (+) or without (-) _int_ercellular wave prop_aga_tion in non-HF (n_LMS_=44; N=11) and HF (n_LMS_=51; N=11). (E) Faction of intercellular waves (from t_ota_l number of Ca^2+^ waves within same LMS) in non-HF (n_LMS_=14) and HF (n_LMS_=28). (F) Fraction of intercellular Ca^2+^ waves with specific direction (end-to-end: non-HF 16/21 waves and HF 45/109 waves) or (side-to-side: non-HF 5/21 waves and _HF_ 64/109 waves)._(G_) Number of cells involved in intercellular propagation in non-HF (n_LMS_=14) and HF (n_LMS_=28).

### Increased Cx43 lateralization in HF

In cardiomyocytes, Cx43 form GJs primarily at the IDs, but a prominent feature of structural heart disease, such as HF, is the redistribution of Cx43 to the lateral cell borders that could underlie intercellular propagation of Ca^2+^ waves^17,41^. To assess Cx43 lateralization, we performed immunofluorescence staining on matched regions of cryopreserved tissues selected from 5 HF and 5 non-HF human samples included in our functional analysis (**Figure 5A**). Cx43 staining colocalized with the cell membrane marker WGA, but not with the ID marker β-catenin, was considered as lateral Cx43. The percentage of Cx43 lateralization was significantly higher in HF tissue, with a mean of 26.9% compared to 18.6% in non-HF (**Figure 5B**). Moreover, cells with Cx43 lateralization appear more in clusters in HF compared to non-HF (**Figure 5C, *Figure S6***), with areas of extracellular matrix (ECM) surrounding these clusters (**Figure 5C, *Figure S7***).

**Figure 5.**
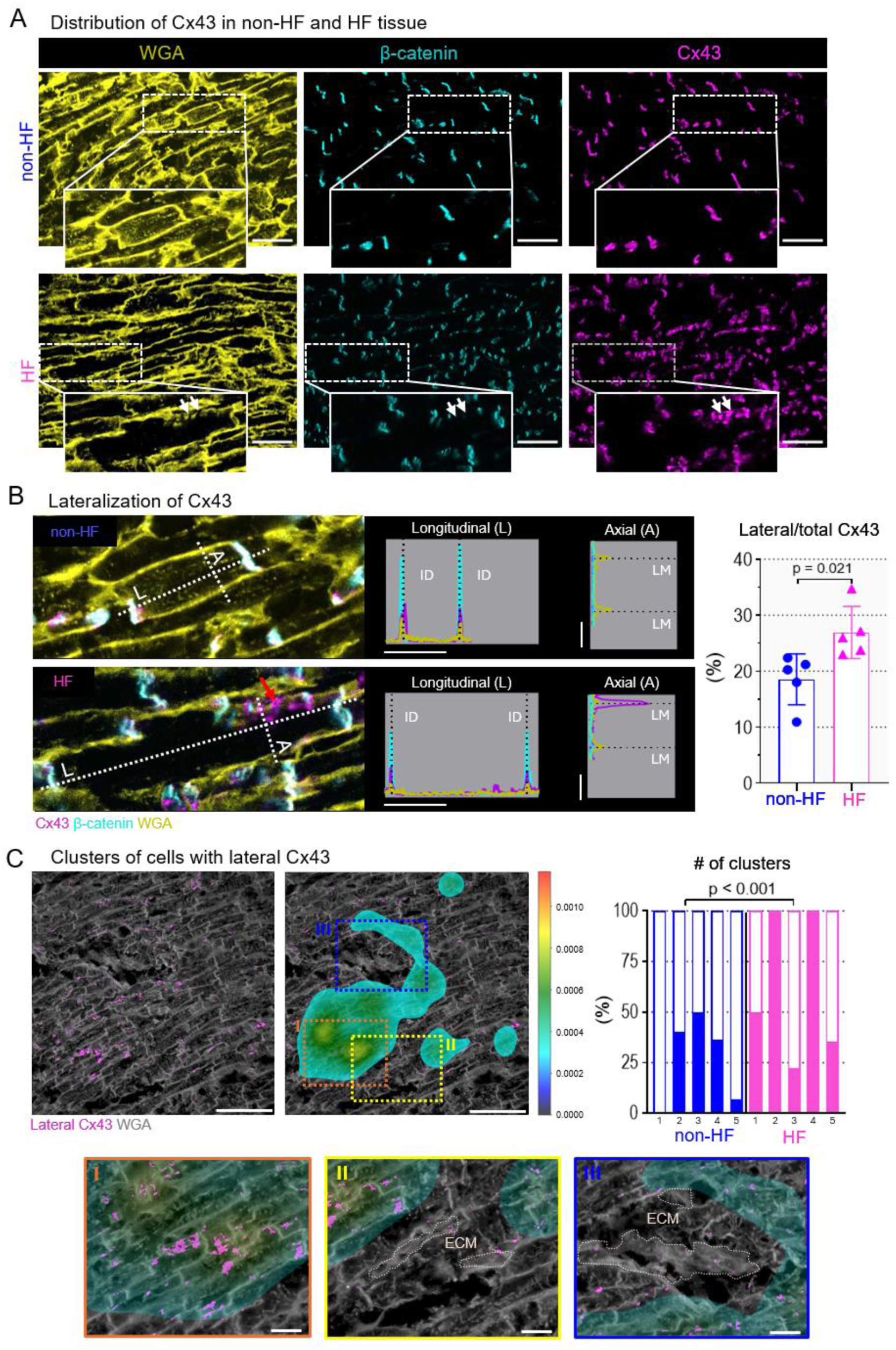
Cx43 lateralization in HF. **(A)** Example images of Cx43 (magenta), β-catenin (cyan) and WGA (yellow) in non-HF and HF tissue sections. Inserts show zoom in and lateral cardiomyocyte membranes (white arrows). Scale bar equals 50 µm. **(B)** Left: Zoom in and merge of 3 channels from panel A. Longitudinal and axial line plots, with distribution of proteins along each axis. The red arrow indicates the distribution of proteins on the lateral membrane. Right: Percentage of lateral Cx43 in non-HF (N=5) and HF (N=5) tissue (ID: intercalated disk, LM: lateral membrane). **(C)** Examples of the organization and quantification of cell clusters with lateral Cx43 (magenta) in tissue. Clusters with lateral Cx43 are shown as a density map ranging from small (black-blue) to large clusters (yellow-orange). The threshold to cluster is set to 0.0003. A zoom in is shown on an example cluster (orange box) and deposits of extracellular matrix (ECM) surrounding these clusters (yellow & blue box). The scale bar equals 100 µm, and 20 µm for zoomed-in areas.

### Increased synchronization of Ca^2+^ waves in HF

Next, we investigated whether the functional and structural alterations observed in HF contributes to tissue-wide synchronization of cells that could initiate arrhythmic activity at tissue level. We distinguished two levels of synchronization resulting in local or global triggered activities (**Figure 6A**). Local triggered activity refers to a group of neighboring cells that have synchronized Ca^2+^ transients, while global triggered activity is the occurrence of LMS-wide synchronized Ca^2+^ transients (**Figure 6A, *Video S7-9***). Overall, triggered activity (both local and global) was observed with a higher incidence in HF compared to non-HF (**Figure 6B**). In more detail, 12.7% of HF LMS (7 out of 55 LMS) exhibited ‘global’ synchronization with a median [interquartile range (IQR)] latency time of 0.93 s [0.83 to 13.76 s], while only 1.8% of non-HF LMS (1 out of 57 LMS) showed this type of synchronization with a latency time of 19.93 s (**Figure 6B-C**). ‘Local’ synchronization was observed in 23.6% of HF LMS (13 out of 55 LMS), compared to 7% of non-HF LMS (**Figure 6B-C**). The latency for these ‘local’ events was comparable between HF (median [IQR]: 0.86 s [0.32 to 2.11 s]) and non-HF (0.55 s [0.30 to 3.76 s]) LMS (**Figure 6C**). The areas from which these events originated consisted of a median of 6 and 3 cells in HF and non-HF LMS, respectively (**Figure 6D**). Importantly, these local triggered activities were able to induce local tissue depolarization, an observation which was also seen for inter-beat Ca^2+^ waves (**Figure 6E**).

**Figure 6.**
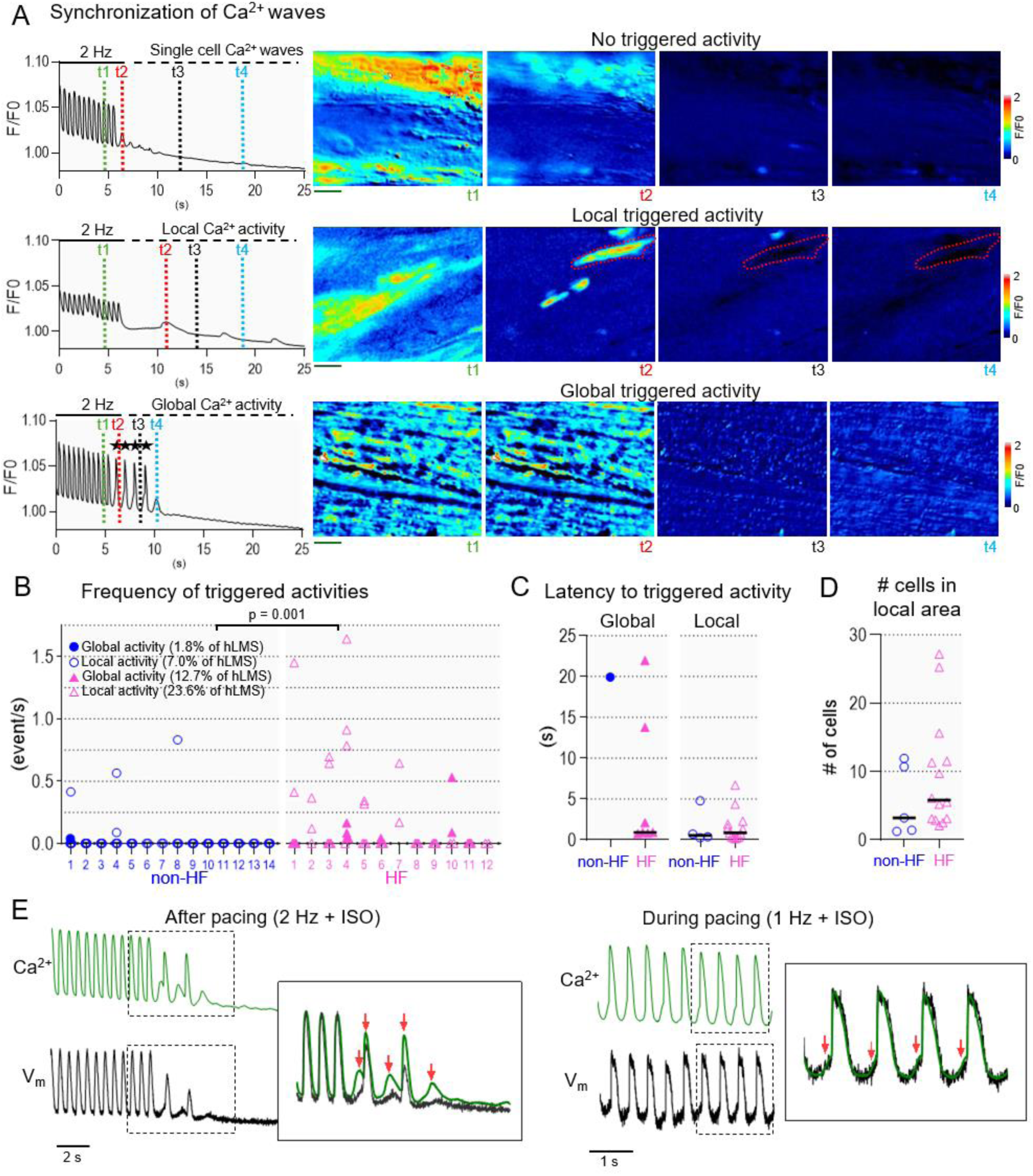
Increased synchronization and triggered activity in HF. **(A)** Example traces and images of different triggered activities after pacing: no (upper), local (middle) or global (bottom) triggered activity. Time frames are indicated with t_x_. **(B)** Frequency of local and global triggered activity in non-HF (N=14) and HF (N=12). **(C)** Latency to triggered activity in non-HF and HF. **(D)** Number of cells in areas with local triggered activity in non-HF and HF. **(E)** Example traces of Ca^2+^ (upper) and voltage (lower) during pacing at 1 Hz in presence of isoproterenol (ISO). Inserts show zoom in, with Ca^2+^-induced changes in voltage (V_m_) (red arrows).

### Ca^2+^ wave dynamics in HF determine the potential for focal excitation

To investigate whether experimentally-observed dynamics of Ca^2+^ waves are sufficient to lead to focal excitations in tissue, we used a recently proposed modeling approach that enables the direct control of the timing and amplitude of waves to be imposed in cell and tissue models^34^. First, cellular responses to spontaneous Ca^2+^ release were studied. For this, individual SRF were imposed in non-HF and HF cell models with a set latency time and gradually increasing the amplitude from very low amplitude events to very large. This allows the amplitude threshold for triggered APs to be extracted (**Figure 7A**). Due to the reduced *I*_K1_ and increased *I*_NaCa_, the HF model exhibited larger DADs than the non-HF model for a given amplitude of Ca^2+^ waves, which manifested as a lower threshold for the triggered APs (N_RyR_ peak of 0.09 compared to 0.21 in non-HF; **Figure 7A**). Furthermore, the emergence of focal excitations was studied in tissue models. Only in the remodeled tissue condition, where intercellular connectivity was reduced, focal excitations were observed. Therefore, all the following data are from the remodeled tissue condition. In both non-HF and HF electrophysiological conditions, the dynamics of Ca^2+^ waves observed in HF led to a larger number (higher probability) of focal excitations. In more detail, in non-HF electrophysiology, no focal excitations were observed when imposing the latency time distributions from non-HF observations, whereas few were when imposing the latency time distributions from HF observations (**Figure 7B, left**). With HF electrophysiology, substantially more focal excitations were observed using the HF latency time distributions than the non-HF ones (**Figure 7B, right**). Two examples of focal excitation are illustrated in **Figure 7C**. One case is shown with Ca^2+^ wave dynamics in non-HF, wherein a focal excitation emerges late after multiple, relatively small DADs, another case shows Ca^2+^ wave dynamics in HF, where large DADs are observed and a focal excitation emerges relatively quickly. Therefore, cellular remodeling, Ca^2+^ wave dynamics differences, and tissue remodeling in HF all combine to promote focal excitations emerging, each contributing to the observed increase in arrhythmia vulnerability.

**Figure 7.**
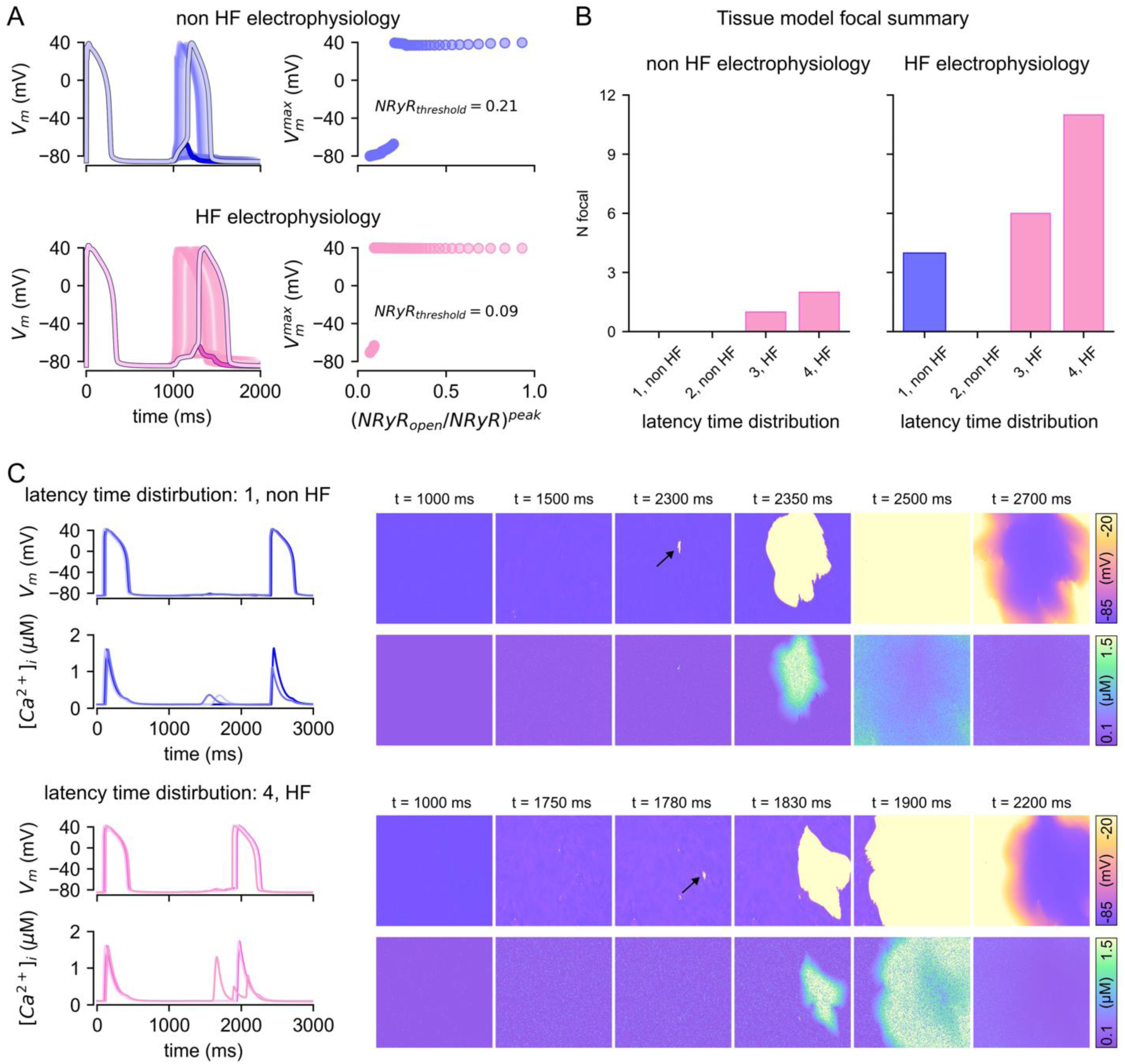
Ca^2+^ waves in HF can induce triggered activity. **(A)** Illustration of the threshold for a triggered action potential (AP) in a single cell. Left panel: Time course of voltage corresponding to a paced excitation and imposition of spontaneous release function (SRF) at gradually increasing amplitude, overlayed, with examples either side of the threshold (largest non-triggered DAD, first triggered AP) highlighted. Right panel: Maximum voltage after paced AP against amplitude of the SRF (measured in peak N_RyR__open/N_RyR_), illustrating where the threshold occurs. **(B)** Summary of total number of focal excitations under each condition. All data are for the remodeled tissue model. **(C)** Illustration of synchronization and focal excitation in different conditions in tissue. Left panels: voltage and Ca^2+^ concentration for three arbitrarily selected nodes in the tissue, right panels: snapshots of the spatial distribution of voltage and Ca^2+^. The voltage color scale is clipped with −20 mV for clarity, though actual maximum voltage exceeds this (around +40 mV).

## DISCUSSION

We have identified several mechanisms at single-cell and cell organizational level that underlie transition from increased cellular spontaneous Ca^2+^ release in HF to tissue-wide triggered activity under conditions of fast-pacing and adrenergic activation. In HF, Ca^2+^ waves occur more frequently early after a paced beat and in close proximity to each other. Moreover, in HF, Ca^2+^ waves propagate faster and are more likely to propagate from its initiating cardiomyocyte to a neighbour. Among these intercellular waves, the transverse direction is more prevalent, together with an increase in lateralized Cx43. These mechanisms together contribute to synchronization and the appearance of clusters of cells with triggered activity, with the potential to induce tissue-wide PVC and initiate arrhythmias.

### Altered dynamics of cellular Ca^2+^ waves in time and space in HF

Even in single cardiomyocytes, single Ca^2+^ waves are rarely sufficient to trigger an AP. In HF, cardiomyocytes have a higher propensity for spontaneous Ca^2+^ release, increasing the likelihood of triggering an AP, through summation of multiple Ca^2+^ waves, combining into larger NCX currents, and more unstable resting membrane potential, activating Na^+^ channels^26,27^. At tissue scale, however, the combined NCX current needs to be sufficiently large to overcome the electrotonic current drain from neighboring cells (source-sink mismatch)^42^. In HF, altered dynamics of spontaneous Ca^2+^ release in time and space overcome this mismatch, as the frequency of Ca^2+^ waves in neighboring cells is high at approximately the same time. We identified several mechanisms that further contribute to these requirements for propagation. Firstly, early onset Ca^2+^ waves occur more frequently in HF. By dissecting Ca^2+^ waves into different time bins, we uncovered that more Ca^2+^ waves occur in the first 2 s post-pacing in HF hearts. It is important to note that when averaging the total number of events over the full duration of recording (i.e., 20 s), the number of Ca^2+^ waves was not different between non-HF and HF tissue. This suggests that in HF, there is an optimal substrate after pacing that rapidly decays. Previous studies have identified a role for CaMKII activation to increase RyR open probability, and in the case of HF, this would be typically enhanced through pacing and adrenergic activation^27,43^. Secondly, early onset Ca^2+^ waves travel faster in HF leading to accumulation of the cytosolic Ca^2+^ load and hence the magnitude of the depolarizing NCX current in each cell. The higher velocity of Ca^2+^ waves reflects a facilitated RyR to RyR propagation, likely attributable to increased RyR sensitivity and/or elevated Ca^2+^ loading at the wave front, and thus a lower SR-Ca^2+^ release threshold^44–47^. In addition to the functional RyR modulation, organization of RyR clusters and structural remodeling in HF myocytes, can modulate RyR function to sustain increased Ca^2+^ wave frequencies and propagation. These changes include altered T-tubular networks^48,49^ and consequent local microdomain signaling near RyRs^26,27^, fragmentation of RyR clusters^50^, increased IP_3_R expression that could facilitate intercluster coupling^26^, and rearrangement of RyR cluster interspacing^51^.

### Altered cell-cell connectivity facilitates propagation of Ca^2+^ waves between cells

We found an increased potential for intercellular propagation of Ca^2+^ waves in HF tissue. Traditionally, cardiomyocyte pairs isolated by incomplete digestion from adult hearts have been the primary model for studying intercellular Ca^2+^ flow^52–54^, focusing mainly on one-dimensional end-to-end propagation, which restricts understanding of cellular anisotropy and radial wave dynamics. Intact cardiac tissue from rodents has also been used to study intercellular Ca^2+^ waves propagation^55–58^. Similar to our data, intercellular propagations are not frequently observed and the path of cell-to-cell propagation involved only few neighboring cardiomyocytes. These studies found intercellular propagation to be slow^55–58^, making it unlikely that cell-to-cell propagation of Ca^2+^ waves is the only source to cause arrhythmias.

Furthermore, in line with other studies^59,60^, we found that the end-to-end propagation of Ca^2+^ waves between cells is less likely. Instead, Ca^2+^ waves propagate more readily in side-by-side cell configuration, which coincides with the increased Cx43 lateralization. Furthermore, a sparse junctional SR at the ID and end-to-end junctions might impede propagation of Ca^2+^ waves in a side-to-side configuration^59,60,61^. Indeed, modeling studies predict that propagation is likely to fail when sarcomere length exceeds the normal value of 1.9 µm, and thus can act as a preventive mechanism against aberrant propagation in longitudinal direction^62^. Other mechanisms, such as increased local buffering near the ID (e.g., presence of mitochondria), inducibility of Ca^2+^ waves in the receiving neighboring myocyte (e.g., RyR sensitivity) and GJ conductivity can further support differences in potential of intercellular propagation.

### Heterogeneity and clusters of focal activity

In HF, cardiomyocytes with early onset Ca^2+^ waves appear in clusters, creating a larger source to overcome the current sink. Since arrhythmia vulnerability in cardiomyopathy stems from cell-cell variabilities of excitability, repolarization, and gene expression^63,64^, cardiomyocyte heterogeneity likely drives the formation of local clusters. In this case, phenotypic heterogeneity leads to variations in Ca^2+^ dynamics, with a higher fraction exhibiting early onset wave activity compared to surrounding cells. Additionally, the synchronization of large clusters is likely determined by the degree of intercellular coupling. As shown in numerous models, in a strongly heterogenous cell population, the intercellular coupling strength influences the synchrony within a system. Additionally, heterogeneously remodeled intercellular coupling can contribute to heterogeneity in other cellular properties, such as AP duration and excitability, enhancing the diverse cellular environment that promotes arrhythmia formation. Modelling studies suggest that a large number of cardiomyocytes are required to induce ectopic activity at the tissue level, ranging from hundreds to millions of cells^32,65^. Multiple local clusters may accumulate or interact to collectively create sufficient source for arrhythmic events. Indeed, 2D simulations based on Ca^2+^-voltage parameters from rabbit cardiomyocytes propose that synchronized Ca^2+^ waves in clusters contribute to focal activity^42^. Both our study and other in silico models predict reduced latency periods of Ca^2+^ waves across different cells in the myocardium with minimal variance, to allow for effective summation of diastolic Ca^2+^ waves within a cluster^42^. In a CPVT study using murine slices with a leaky RyR mutant, frequent diastolic Ca^2+^ waves were shown that directly and adversely engage surrounding excitable cells. A similarly short latency period of ∼2 s as the drive for focal arrhythmias was found important^31^. Furthermore, GJ coupling between cardiomyocytes may have a dual-acting effect on the source-sink balance in focal activity formation. In vulnerable clusters, depolarizing currents from ‘source’ cells can flow through GJs, driving Ca^2+^ leak in adjacent cells and amplifying synchronous waves. At the whole heart level, these local regions are poorly coupled to the rest of the myocardium as a result of heterogenous GJ remodeling or as shown here by ECM deposits in HF development^66^. Additionally, GJs at the ID are globally reduced in the human HF tissue which can engage in disrupting the normal source-sink mismatch.

### Implications for arrhythmia mechanisms in HF

Our experimental findings demonstrate that in HF hearts, diastolic Ca^2+^ waves occur with early onset are more prevalent in clusters of adjacent cardiomyocytes. The pivotal role of this synchronous activity in triggering focal excitation was identified through 2D simulations, considering failing cardiomyocyte electrophysiology, reduced cellular connectivity in HF tissue, and the experimentally attained latency distribution. The short coupling interval of early onset Ca^2+^ waves aligns with diastole in the working myocardium, where we similarly observed an increased number of Ca^2+^ waves in HF hearts. This short coupling interval presents an increased risk of propagation failure, as in this time window, ectopic foci are more likely to cause wave break and fibrillation. Fibrosis can exacerbate this risk by slowing the excitation wavefront, creating conduction block, and supporting re-entrant circuits, especially in ischemic cardiomyopathy, where local fibrotic areas near the infarct are more vulnerable to induce ectopic activity^67,68^. In contrast, other cardiomyopathies, characterized by diffuse fibrosis and homogeneously distributed remodeling across the myocardium, remain less well studied. Nevertheless, our findings point to a common arrhythmogenic mechanism involving cell clusters of early-onset Ca^2+^ waves in different HF etiologies.

### Strengths and Limitations

The study of LMS allows linking cellular events to tissue properties and uncovers a level of heterogeneity relevant to in vivo arrhythmogenesis. We could capture critical temporal dynamics that facilitate focal and propagated events, adding information beyond studies of RyR function in isolated cells, which often average Ca^2+^ events over time (>10 s). Study-limitations include myofilament-induced Ca^2+^ waves upon stretch, which were unassessed due to inhibition of contraction. Furthermore, the temporal resolution of 200 Hz restricted resolving direct causal links between intercellular Ca^2+^ waves and Cx43 remodeling or measuring GJ coupling strength globally vs. locally.

### Conclusions

In human HF LMS, we found a higher incidence of early onset Ca^2+^ waves with increased synchrony from clusters of nearby cells. In conjunction with altered intercellular connectivity, these localized arrhythmogenic waves can overcome the current sink to depolarize the myocardium, leading to focal activities in HF.

## ABBREVIATIONS

AP: Action potential
Cx43: Connexin43
DAD: Delayed afterdepolarization
HF: Heart failure
ID: Intercalated disc
ISO: Isoproterenol
LMS: Living myocardial slices
NCX: Na^+^/Ca^2+^ exchanger
Non-HF: Non-failing heart
PVC: Premature ventricular complex
RyR: Ryanodine receptor
SD: Standard deviation
SRF: Spontaneous release function

## Acknowledgements

We would like to thank the division of Cardiac Surgery in the Department of Cardiovascular Sciences at KU Leuven and the UZ Leuven transplant team for help in providing human explanted hearts, Roxane Menten for coordination and members of the Laboratory of Experimental Cardiology for biobanking of human tissue samples, and Geert Molenberghs for expert advice on the statistical analyses.

## Supplemental Material

Extended Materials and Methods

Figure S1-S7

Table S1-S2 Video S1-S9

Supplemental References Major Resources Table

